# High-throughput TCRB enrichment sequencing of human cord blood exhibited a distinct fetal T cell repertoire in the third trimester of pregnancy

**DOI:** 10.1101/2022.09.08.506871

**Authors:** Yan Dong, Wei Chen, Jinmin Wang, Xiaolei Wu, Yangyu Zhao, Yuhang Cai, Yingxin Han, Yuqi Wang, Hongmei Li, Jie Qiao, Yuan Wei

## Abstract

**Study question:** What are the molecular characteristics during the maturation process of the human fetal immune system in the third trimester of pregnancy?

**Summary answer:** Both the diversity and length of complementarity determining region 3 (CDR3s) in the fetal TCRB repertoire were less than those of adult CDR3s, and the fetal CDR3 length increased with gestation weeks in late pregnancy.

**What is known already:** The adaptive immune system recognizes various pathogens based on a large repertoire of T-cell receptors (TCR repertoire), but the maturation dynamics of the fetal TCR repertoire in the third trimester are largely unknown. The CDR3is the most diversified segment in the T-cell receptor β chain (TCRB) that binds and recognizes the antigen.

**Study design, size, and duration:** This was a basic research to assess the composing characteristics of TCRBs in core blood and the dynamic pattern with fetal development in the third trimester of pregnancy.

**Participants/materials, setting methods:** High-throughput TCRB-enrichment sequencing was utilized to characterize the TCRB repertoire of cord blood at 24~38 weeks of gestational age (WGA) with nonpreterm fetuses and to investigate their difference compared with that of adult peripheral blood.

**Main results and the role of chance:** Compared to the adult control, the fetal TCRB repertoire had a 4.8-fold lower number of unique CDR3s, a comparable Shannon diversity index (*p*=0.7387), a lower mean top clone rate (*p* < 0.001) and a constrictive top 1000 unique clone rates. Although all kinds of TCRBV and TCRBJ genes present in adult CDR3s were identified in fetuses, nearly half of these fragments showed a significant difference in usage. Moreover, the fetal TCRB repertoire held a shorter CDR3 length, and the CDR3 length showed a progressive increase with fetal development. Jensen–Shannon (JS) divergences of TCRBV and TCRBJ gene usage in dizygotic twins were much lower than those in unrelated pairs. In the parental-fetal pair, JS divergence of TCRBV gene usage was not obviously different, while that of TCRBJ gene usage was only slightly lower.

**Limitations, reasons for caution:** The sample size is limited due to the limited accessibility to cord blood in late pregnancy with healthy nonpreterm fetuses.

**Wider implications of the findings:** Our findings reveal the unique properties of fetal TCRB repertoires in the third trimester, fill the gap in our understanding of the maturation process of prenatal fatal immunity, and deepen our understanding of the immunologically relevant problems in neonates.

**Study funding/competing interest(s):** This work was supported by the National Natural Science Foundation of China (82171661) and Tianjin Municipal Science and Technology Special Funds for Enterprise Development (NO. 14ZXLJSY00320). The authors declare that they have no competing interests.

## Introduction

The ability of the adaptive immune system to recognize a wide variety of pathogens relies on a large repertoire of unique T-cell receptors (TCRs) (Qi et al., 2014). The TCR that mediates the response to antigenic peptide major histocompatibility complex (MHC) plays a major role in controlling the selection, function and activation of T cells (Han et al., 2014, Woodsworth et al., 2013). The T-cell receptor β chain (TCRB) consists of V, D, J, and C gene segments. Complementarity determining region 3 (CDR3) generated by V-(D)-J recombination is the most diversified segment to bind and recognize the antigen. The diversity of the TCR repertoire mainly results from rearrangements of various gene segments, imprecise joining, the nibbling of germline nucleotides and the addition of N- and P-residues at the V-D-J junction sites (Casrouge et al., 2000, Garderet et al., 1998). The unique TCRB sequences of young adults have reached 100 million for a minimal estimate (Qi, Liu, Cheng, Glanville, Zhang, Lee, Olshen, Weyand, Boyd and Goronzy, 2014).

Fetal immune system undergoes a developing process during the whole gestation (Park et al., 2020). Researchers have made great progress in profiling the ontogeny of the human immune system and have depicted the developmental dynamics of immunity in various fetal organs and blood circulation system during pregnancy, especially in the first and second trimesters of gestation(Feyaerts et al., 2022, Park, Jardine, Gottgens, Teichmann and Haniffa, 2020, Rechavi et al., 2015, Suo et al., 2022). Putative prothymocytes can be detected from 7 weeks, and T-cell precursors derived from the fetal liver then seed the thymus at 8-9 weeks(Holt and Jones, 2000). From 9.5 weeks, TCRB-positive cells emerge and increase and form over 90% of the CD7-positive population until birth (Campana et al., 1989). Mature T cells can be observed in the circulatory system at 15-16 weeks(Holt and Jones, 2000). Erez Rechavi and his colleagues collected fetal blood under fetal reduction and cardiocentesis between 12 and 26 WGAs and revealed reduced diversity and uneven representation of clonotypes in the fetal T cell repertoire at early gestational age but a progressive increase in diversity and evenness with gestation weeks(Rechavi, Lev, Lee, Simon, Yinon, Lipitz, Amariglio, Weisz, Notarangelo and Somech, 2015). Meanwhile, although TCRB repertoire diversity at later second and early third trimesters (22 to 26 WGA) was comparable with that in healthy infants, the average CDR-B3 length increased with gestation weeks and was significantly shorter than that in healthy infants(Rechavi, Lev, Lee, Simon, Yinon, Lipitz, Amariglio, Weisz, Notarangelo and Somech, 2015). The cord blood (CB) collected at delivery has been thoroughly studied in recent decades (Basha et al., 2014, Britanova et al., 2016, Fadel and Sarzotti, 2000, Guo et al., 2016, Harris et al., 1992, Zhao et al., 2019). It has been reported that the length of the third complementarity-determining region of the heavy chain (HCDR3) of cord blood progressively increases during the third trimester(Schroeder et al., 2001). Brian L. Le and his colleagues compared the TCRB repertoires of cord blood obtained from term and preterm deliveries (32.4 to 35.2 WGA) and showed shorter CDR3s and skewed usage of the V, D, J genes in preterm infants(Le et al., 2021). These studies suggested the ongoing maturation of fetal T cell-media adaptive immunity in the third trimester. However, emollient direct evidences was still absent, and the detailed characteristics of the T cell repertoire of healthy nonpreterm fetuses during this period remain largely unknown due to limited accessibility to samples.

Multitudinous studies about immune repertoire were targeted on a limited number of sequences. (Schelonka et al., 1998, Schroeder, Zhang and Philips, 2001, Souto-Carneiro et al., 2005). The high-throughput sequencing of genes coding immune receptors enables researchers to perform a comprehensive assessment of the immune repertoire (Ghraichy et al., 2020, Guo, Wang, Cao, Yang, Liu, An, Cai, Du, Wang, Qiu, Peng, Han, Ni, Tan, Jin, Yu, Wang, Wang, Wang and Ma, 2016, Le, Sper, Nielsen, Pineda, Nguyen, Lee, Boyd, MacKenzie and Sirota, 2021). Although a previous study tried to apply high-throughput TRB/IGH enrichment sequencing to decode the characteristics of T and B-cell repertoires of fetal blood at different stages of pregnancy, the sample size was small, with only one sample in each of the four stages no later than 26 WGA(Rechavi, Lev, Lee, Simon, Yinon, Lipitz, Amariglio, Weisz, Notarangelo and Somech, 2015). Here, we collected cord blood samples in the third trimester of pregnancy by cordocentesis and utilized high-throughput TCRB enrichment sequencing to comprehensively profile the unique characteristics of the fetal TCRB repertoire during this period, as well as their differences compared with those of adults.

## Materials and Methods

### Ethical approval

This study was conducted with ethical approval obtained from the Reproductive Study Ethics Committee of Peking University Third Hospital. (approved protocol no. 2013SZ025). All participants provided written informed consent for participation.

### Sample collection and treatment

Fetuses who were with or suspected of conditions known to have an effect on immune development or pregnant women who had ever suffered from immune-related diseases or received any immune-related therapy in the past months were excluded from this study. Adult controls were volunteers in BGI who had also received written informed consent.

Thirteen pregnant women at an average age of 30.4 were recruited by PUTH, and some of them underwent cordocentesis for a higher risk for fetal aneuploidy or fetal ultrasound abnormality in pregnancy; nevertheless, the infants were phenotypically normal at delivery. A total of 14 cord blood samples were collected at 24 to 38 WGA (median gestational age, 28 weeks and 6 days) by PUTH, while the pregnant women were at delivery or underwent a cordocentesis. No immunodeficiency was documented in any of the families. In detail, in these fourteen cases, three samples were obtained after full-term born delivery (CB7, CB13, CB14), including a dizygotic twin pair (CB13 and CB14), while the remaining eleven samples were collected during cordocentesis (details are provided in **Table 1**).

Each sample with 1 mL to 2 mL of cord blood was stored in an EDTA-containing tube. Peripheral blood mononuclear cells (PBMCs) were isolated in two hours by using Ficoll Paque Plus (17-1440-02, GE Healthcare), deposited in RNAlater (Am7021#RNAlater Soln. INVITROGEN), and stored at −80°C. Equivalent peripheral blood samples from parents of CB5 and fourteen healthy adult controls (average age 31.2 ± 2.8 years; no pregnant women) were handled in the same way.

### Library construction

Total RNA was extracted from PBMCs according to the manufacturer’s protocol (80204, AllPrep DNA/RNA Mini Kit, QIAgen) and evaluated using an Agilent 2100 Bioanalyzer (5067-1513, Agilent RNA 6000 Pico Kit). After DNase I (M0303S, NEB)treatment, reverse transcription was performed using 200 ng of total RNA mixed with 20 ng random hexamer primer, 1 μL dNTP mix (10 mM each, N201L, ENZYMATICS), and DEPC-treated water(AM9915G, AMBION) to make a total volume of 12 μL. The samples were incubated for 5 minutes at 65°C and quickly chilled on ice. The contents of the tube were collected by brief centrifugation, and 4 μL 5X first-Strand Buffer, 1 μL DTT (0.1 M), and 1 μL RNaseOUT (40 units/μL,10777019, INVITROGEN) were added. The contents were gently mixed and then incubated at 25°C for 2 minutes. After the addition and mixing of 1 μl SuperScript™ II RT(18080-044, INVITROGEN), RT was performed at 42°C for 50 minutes before heat inactivation at 70°C for 15 minutes. Subsequently, 5 μl of the reverse transcribed products was amplified with the Vβ forward primers and Jβ reverse primer set pools (0.2 μM each) using a Multiplex PCR Kit (206143, QIAGEN, Germany). The following PCR program was used: 1 cycle of 95°C for 15 minutes, 25 cycles of denaturation at 94°C for 30 s, annealing at 60°C for 90 s, and extension for 30 s at 72°C. The last step was final extension for 5 minutes at 72°C and then cooling to 12°C. Size selection was also used for purification of 100 bp-200 bp PCR products by QIAquick Gel Extracton (28706, QIAGEN, Germany).

DNA library preparation followed the manufacturer’s instructions (PE-402-4001, PE HiSeq 2500 Flow Cell, Illumina) as described previously (Wang et al., 2008). We used the same workflow as described elsewhere to perform cluster generation, template hybridization, isothermal amplification, linearization, blocking and denaturization and hybridization of the sequencing primers. Paired-end sequencing of samples was carried out with a read length of 100 bp using the Illumina Hiseq2500 platform. In total, we obtained an average of 8.85 M raw reads for each sample.

### Analysis of Illumina sequence data

We first merged the high-quality paired reads using COPE and FqMerger (BGI) and designated the results as contigs. Then, we seted a reference directory constituted by sets of sequences that contain the human (Homo sapiens) TCRB V-REGION, D-REGION and J-REGION alleles (http://www.imgt.org/). The TCR CDR3 region as defined by the International ImMunoGeneTics (IMGT) collaboration begins with the second conserved cysteine encoded by the 3 portion of the V gene segment and ends with the conserved phenylalanine encoded by the 5 portion of the J gene segment. The TCRB CDR3 regions were identified within the sequencing reads according to the definition established by the IMGT collaboration (Giudicelli and Lefranc, 2011). Finally, we obtained an average of 8.41 M reads for each sample. TPM normalization was used before the comparison between the samples. More details in the analysis of Illumina sequence data have been reported in the literature (Han et al., 2015, Zhang et al., 2015).

### Statistical methods

Statistical analysis was performed, and graphs were made using the R statistical programming language (version 2.15.3) and package *ggplot2* (version 0.9.3.1). The *P value* was calculated using the Mann–Whitney test or t test, and a single asterisk (*) indicates *p ≤ 0.05*, double asterisks (**) indicate *p ≤ 0.01*, and triple asterisks (***) indicate *p ≤ 0.001*. *p <0.05* was regarded as significant.

Jensen–Shannon divergence (JS divergence) was defined as follows:

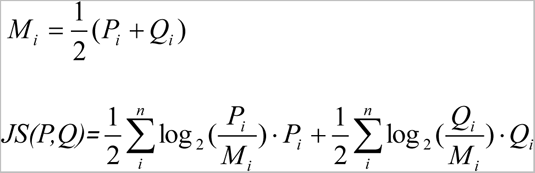

*P_i_*, *Q_i_* is the V or J gene usage of individuals.

## Results

### Reduced diversity and even distribution of the TCRB repertoire in fetuses compared with adults

As each unique CDR3 sequence denoted a T-cell clone (Thapa et al., 2015), we assessed the number of unique CDR3 by uniform random sampling of 4 million reads from each sample in each group to compare the diversity of the TCRB repertoire between fetuses and adults. We found that the number of unique CDR3s in the fetus group was 4.8-fold lower than that in the adult group [n ± SD of 46625.57±21889.59 vs 222486.8 ±76738.86, p < 0.001] (**Fig. 1A**), indicating that the TCRB repertoire of the fetus had a reduced diversity.

**Figure 1.**
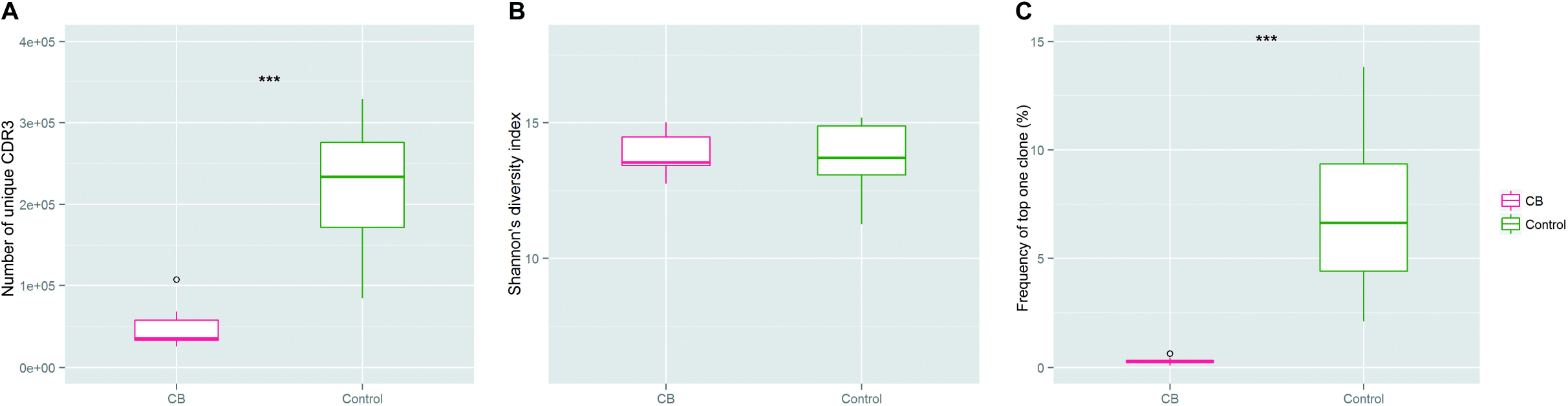
Diversity of the TCRB repertoire in cord blood and adult peripheral blood. (A) Variance analysis of the number of unique CDR3 clones in cord blood and adult peripheral blood TCRB repertoires. The unique CDR3 clone number was calculated using 4 million random reads for normalization (*p<0.001).* (B) Variance analysis of Shannon’s diversity index in cord blood and adult peripheral blood TCRB repertoires (*p*=0.7387). (C) Variance analysis of the top one clone rate in cord blood and adult peripheral blood TCRB repertoire (*p<0.001). *p<0.05, **p<0.01, ***p<0.001* according to two-tailed *t* test.

Shannon entropy was one of the most widely used parameters to assess the immune repertoire diversity, as it integrated not only the number of unique CDR3s but also the relative proportion of each unique CDR3(Six et al., 2013); the higher the Shannon’s diversity index was, the more equal the distribution of each unique CDR3. As shown in **Fig. 1B**, the Shannon’s diversity index of the fetus group and adult group was comparable (p=0.7387). The combination of the reduced diversity and comparable Shannon’s diversity index suggested that the distribution of unique CDR3 in the fetus was more even. As the frequencies of dominant clones had a significant impact on the overall distribution of clones, we compared the top clone rate in each group. The mean value of the top one clone rate in the fetus group was 0.28%, which is significantly lower than the 7.16% in the adult group (p < 0.001) (**Fig. 1C**). Moreover, to assess whether the top one clone was just an exception, we further analysed the top 1000 unique clone rates. According to the results, the rates of all these unique clones were in a narrower range (0.008-0.629%) in the fetus than those in adults (range 0.004-13.82%) (**Fig. 2**), indicating a more even distribution of clones in the fetal TCRB repertoire. Furthermore, we classified 14 fetuses into three teams (T), T1 (24-27 WGA), T2 (32-33 WGA) and T3 (37-38 WGA), and compared the VJ pairing pattern (3D format) between the representative fetus selected randomly from each team and three control adults. The results confirmed the more even distribution in the fetal TCRB repertoire than in the adult TCRB repertoire (**Fig S1**).

**Figure 2.**
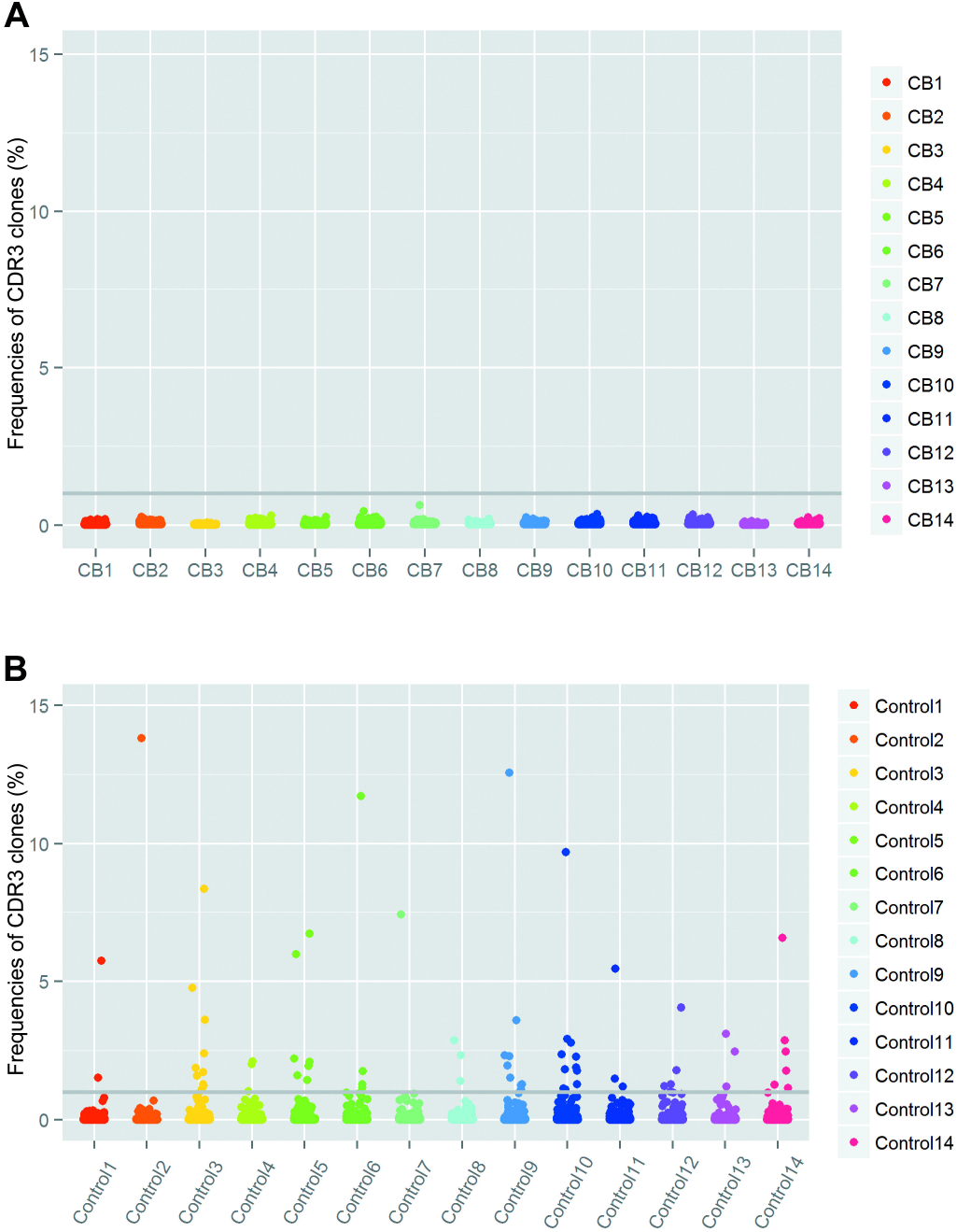
The frequency of the top 1000 CDR3 clones. (**A-B**) The frequency of each clone was determined by calculating the number of reads for each clone divided by the total number of filtered sequencing reads in the sample. The scale of the y-axis was adjusted to an equivalent percentage. Each dot represents a unique clone, and the gray line represents a frequency of 1%. (**A**) Fourteen CB samples. (**B**) Fourteen adult controls.

### Divergent TCRBV and TCRBJ gene usage of CDR3 in fetus and adult

As TCRB CDR3 in the fetus demonstrated a reduced diversity and a more even distribution of unique clones, we assessed the preferential usage of certain V gene and J gene segments by calculating the proportion of sequences belonging to a specific V gene and J gene family. According to the results, all 48 TCRBV genes and 13 TCRBJ genes presented in adult CDR3 were identified in fetuses, while the frequencies of both TCRBV genes and TCRBJ gene usages were apparently different in the fetus and adult groups (**Fig. 3**). Nearly half (25/48) of the TCRBV genes showed a significant divergence (*p* < 0.001) between the two groups, among which 15 genes were significantly overrepresented, and 10 genes were underrepresented. TCRBV genes with higher usage in adults (n=12, frequency > 0.025%) were also used frequently in the fetus, despite a lower frequency. Similarly, a comparable proportion (6/13) of the TCRBJ gene segments showed a significant divergence (*p* < 0.001), apart from TCRBJ1-4 and TCRBJ2-4, all of which were overrepresented. No preferential usages of the D-proximal or D-distal TCRBV gene family and TCRBJ gene family were observed in our study.

**Figure 3.**
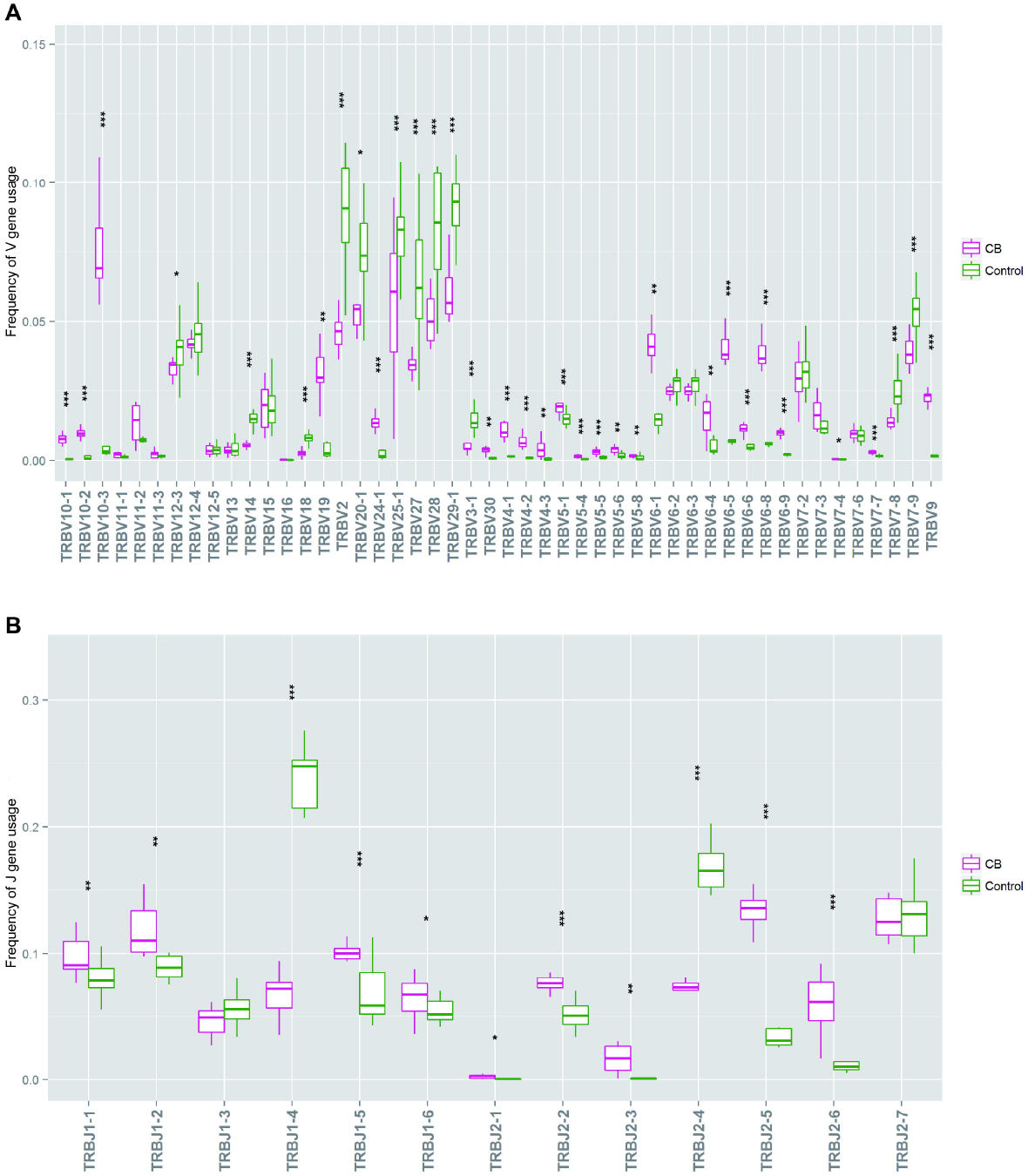
Usage profile of TCRBV and TCRBJ gene families. (**A-B**) Usage profiles of the TCRBV and TCRBJ gene families were assessed by calculating the proportion of sequences belonging to a specific V gene and J gene family (Mann–Whitney test). (**A**) Variance analysis of TCRBV gene usage in the CB and adult control groups. (**B**) Variance analysis of TCRBJ gene usage in the CB and adult control groups.

### Shorter CDR3 length in the fetus compared with the adult

The length of the CDR3 region had a major effect on the three-dimensional structure of the CDR3 loop and therefore antigen binding specificity(Manfras et al., 1999). According to the analysis of CDR3 length of expressed TCRB sequences, we found that fetuses hold a shorter CDR3s compared with adult controls (**Fig. 4A**), and the average length was 11.41 ± 0.33 nt and 12.08 ± 0.20 nt, respectively.

**Figure 4.**
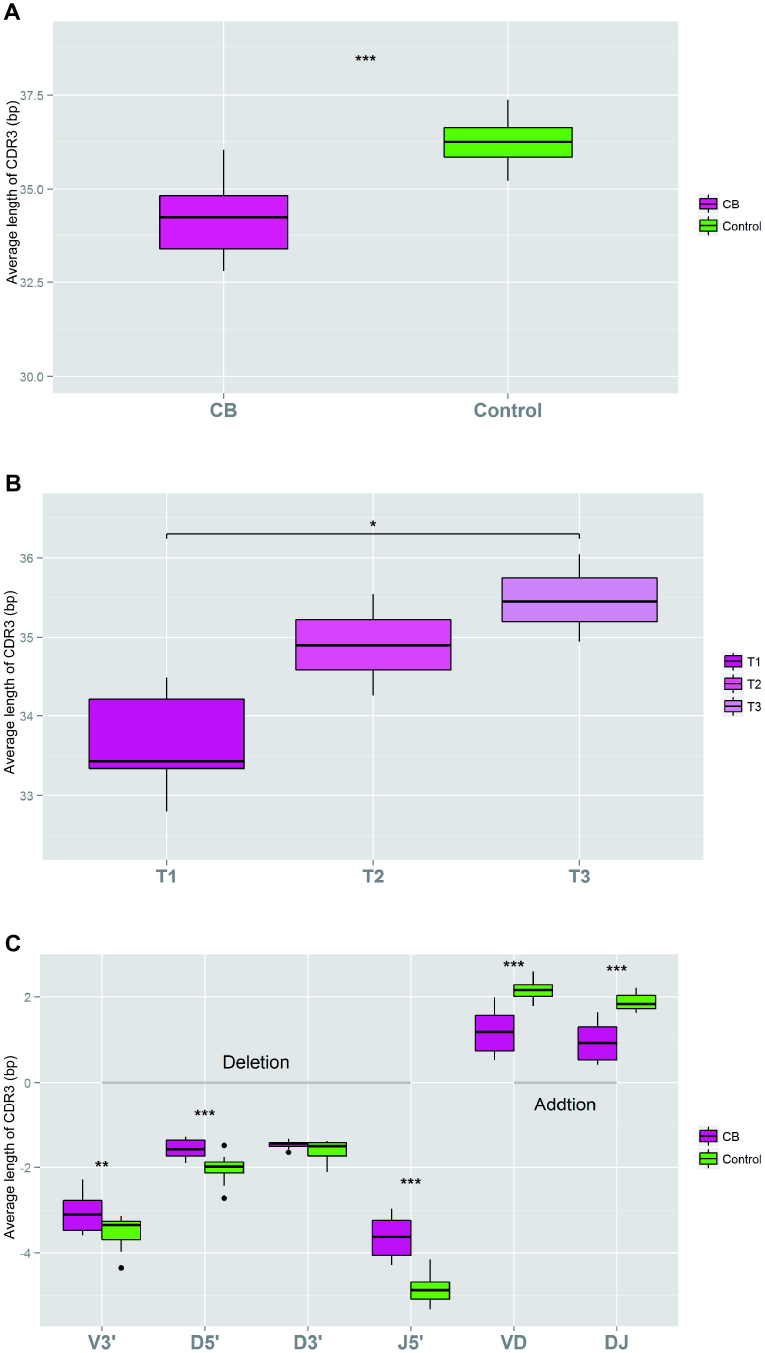
The average CDR3 length in the TCRB repertoire in the CB and adult control groups. (**A**) Variance analysis of the average length of CDR3 in nucleotides (bp) (*p*<0.001, unpaired two-tailed t test). (**B**) Variance analysis of the average length of three teams, T1 (24-27 WGA), T2 (32-33 WGA), and T3 (37-38 WGA). (**C**) Variance analysis of the average length of different components of CDR3. V3’, D5’, D3’, and J5’ represent the nucleotide deletion at the 3’ end of TCRBV, the 5’ end of TCRBD, the 3’ end of TCRBD and the 5’ end of TCRBJ, respectively; VD and DJ represent the nucleotide addition at the TCRBV-TCRBD junction and the TCRBD-TCRBJ junction, respectively.

As significant differences in CDR3 length existed between the adult and fetus groups, we wondered whether there were some differences along with gestational ages. The mean CDR3 lengths of T1, T2 and T3 were 11.20 nt, 11.59 nt and 11.72 nt (**Fig. 4B**), respectively, showing a progressive increase in length along with fetal development.

To find out the probable reasons behind the length differences, we analysed the deletion and addition of nucleotides during the formation of the junctions between V, D and J gene segments (**Fig. 4C**). The mean value of nucleotide deletions in the fetus was 9.73 ± 1.10 nt, while in adults, the value was 11.98 ± 0.47 nt. The number of nucleotide deletions from the 3’ end of TCRBV, the 5’ and 3’ ends of TCRBD, and the 5’ end of TCRBJ were further examined. We found that the fewer deletions from the 5’ ends of TCRBD and the 5’ ends of TCRBJ were mainly responsible for the fewer total deletions (*p* < 0.001). The mean value of nucleotide additions in the fetus sample was 2.14 ± 0.93 nt, while for adult controls, the mean number was 4.03 ± 0.40 nt. Slightly more nucleotides were added at the VD junction than at the DJ junction in both the fetus and adult. Ultimately, taking two factors into account, much fewer nucleotide additions resulted in a shorter net median CDR3 length in the fetus sample.

### Relationship of TCRB repertoires in a pedigree

As genetic factors had an important impact on the initial recombination and selection in the thymus (Zvyagin et al., 2014), we wondered whether there were some potential relationships of TCRB repertoires in a pedigree. We used cord blood samples from a dizygotic twin (CB13 and CB14) and peripheral blood samples from parents of CB5 and quantified the similarity between the TCRBV gene and TCRBJ gene usage in the related and unrelated pairs by Jensen–Shannon (JS) divergence. Lower JS divergence indicated more similar TCRBV or TCRBJ gene distributions. We calculated the JS divergence of the TCRBV gene and TCRBJ gene usage in TCRB repertoires in any possible pairs formed by ten individuals, including a dizygotic twin, a parents-fetus pair, and six unrelated fetuses. As shown in **Fig. 5**, both TCRBV gene and TCRBJ gene usage for TCRB clones were more similar in the dizygotic twin (CB13 and CB14) compared with other individuals, with ~10 times and ~50 times lower JS divergence, respectively (**Fig. 5**, bars 3, 4; **Fig. S2**). However, in the parental-fetal pair (CB5), the JS divergence of TCRBV gene usage was not obviously different from the unrelated divergence, while a slightly lower JS divergence was observed in TCRBJ gene usages (**Fig 5**, bars 6, 7, 8). Overall, the JS divergence of TCRBV or TCRBJ among fetuses was much closer, whereas it was divergent between adults and fetuses.

**Figure 5.**
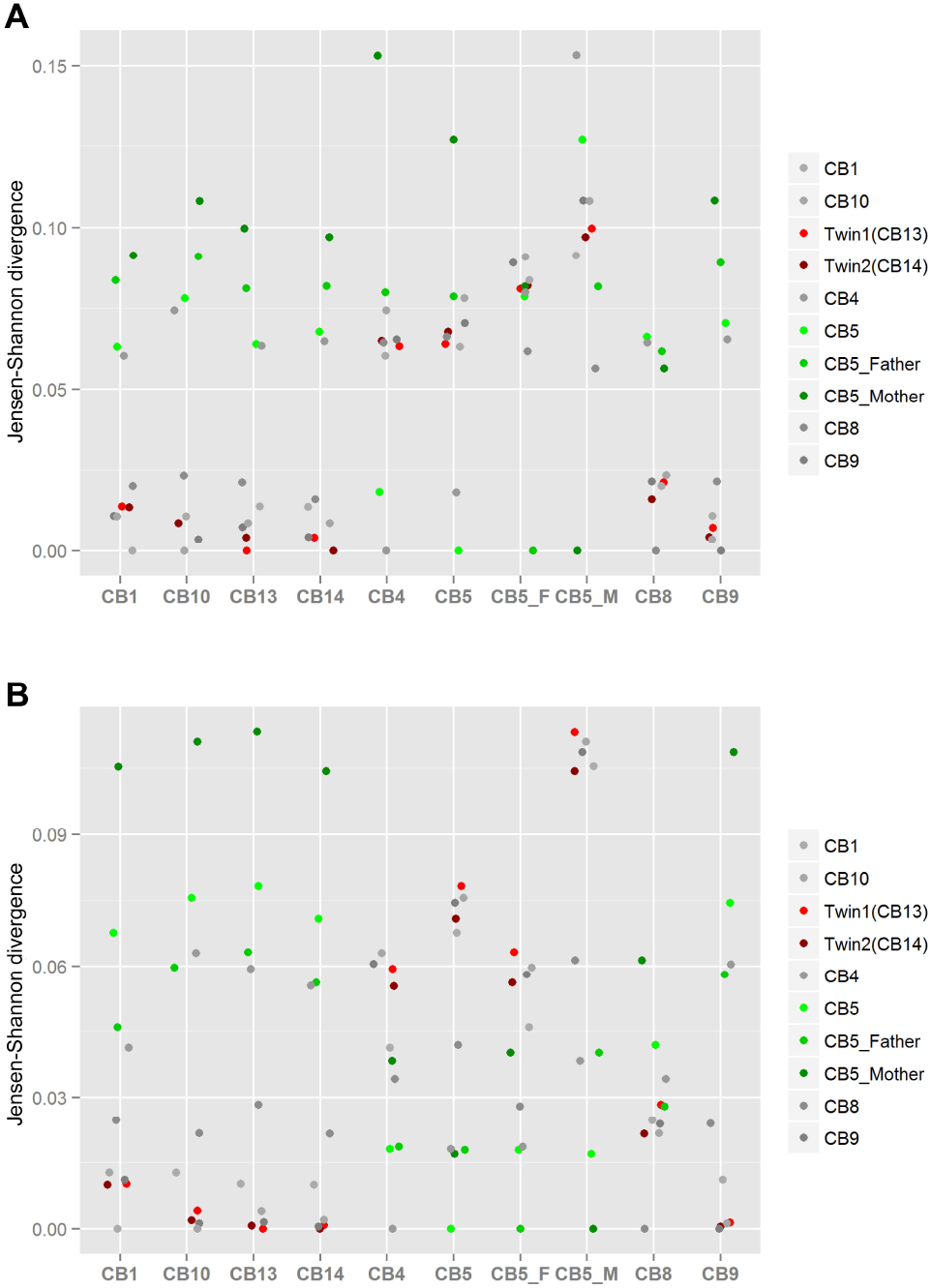
JS divergence for TCRBV and TCRBJ gene usage. (**A-B**) Jensen–Shannon (JS) divergence was used to quantify the similarity between the TCRBV gene and TCRBJ gene usage in each related and unrelated pair. Ten samples were selected, including a pair of dizygotic twin fetuses (red dots), a parental-fetal pair (father, mother and fetus; green dots), and unrelated fetuses (gray dots). (**A**) JS divergence for TCRBV gene usage of in-frame clones. (**B**) JS divergence for TCRBJ gene usage of in-frame clones.

## Discussion

The diversity of TCRB repertoires of different age groups, including neonates, children, adults, and elderly individuals, has been previously published in many works (Basha, Surendran and Pichichero, 2014, Britanova et al., 2014, Garcia et al., 2000, Goronzy and Weyand, 2005, Qi, Liu, Cheng, Glanville, Zhang, Lee, Olshen, Weyand, Boyd and Goronzy, 2014). The TCRB repertoires underwent progressive maturation starting in early gestation (Park, Jardine, Gottgens, Teichmann and Haniffa, 2020, Rechavi, Lev, Lee, Simon, Yinon, Lipitz, Amariglio, Weisz, Notarangelo and Somech, 2015), the profiles of which were relatively less reported due to the difficulties in sample acquisition (Rechavi, Lev, Lee, Simon, Yinon, Lipitz, Amariglio, Weisz, Notarangelo and Somech, 2015). Cord blood samples could be used to characterize the initial frequency and clonal diversity of naive T-cell populations in the fetus, as they emerge from the human thymus and likely before being exposed to any foreign antigens(Moon and Jenkins, 2015). In this study, we systematically assessed the TCRB repertoire of cord blood obtained from 23 WGA to 38 WGA and revealed significant difference between fetus and adult control in the diversity, V gene and J gene usage and CDR3 length. Additionally, we analysed the properties of TCRB repertoires in a pedigree using a dizygotic twin and parental-fetal pair, which had not been reported earlier.

It has been reported that the TRB repertoire of fetal blood in the first trimester is more restricted than that in the late second and early third trimesters (Rechavi, Lev, Lee, Simon, Yinon, Lipitz, Amariglio, Weisz, Notarangelo and Somech, 2015). Our results further demonstrated the reduced diversity of TCRB repertoires in cord blood in the third trimester compared with that of adult control, which was indicated by the 4.8 times lower average number of unique fetal CDR3. The comparable Shannon’s diversity indexes between fetus and adult control and the lower rates of the top one clone and the top 1000 unique clones in fetus together deduced the more even distribution of the clones in the fetal TCRB repertoire. Meanwhile, both our results and another study revealed that the TCR repertoire of CB at delivery was also less complex than that of adult blood (Alfani et al., 2000). The lower diversity and more even distribution profile might reflect the naive nature of T lymphocyte cells in the fetus, indicating that these T cells in CB had not been previously exposed to a high level of antigenic stimulation, while in adults, dominant clonal expansions could generally be considered the hallmark of the antigenic stimuli received throughout life(Garderet, Dulphy, Douay, Chalumeau, Schaeffer, Zilber, Lim, Even, Mooney, Gelin, Gluckman, Charron and Toubert, 1998).

In terms of the CDR3 sequence, diversity is generated centrally through the assembly of V, D and J gene segments and nucleotide trimming plus addition at the junctional region(Le, Sper, Nielsen, Pineda, Nguyen, Lee, Boyd, MacKenzie and Sirota, 2021, Schatz and Ji, 2011). All TCRBV and TCRBJ genes in adults have been used in human fetal life, while each frequency was apparently different. The different usage in the V, D and J gene segments of TCRs compared to that of adults was also mentioned in other studies about the TCR repertoire of cord blood(Guo, Wang, Cao, Yang, Liu, An, Cai, Du, Wang, Qiu, Peng, Han, Ni, Tan, Jin, Yu, Wang, Wang, Wang and Ma, 2016, Rechavi, Lev, Lee, Simon, Yinon, Lipitz, Amariglio, Weisz, Notarangelo and Somech, 2015). Meanwhile, as described in previous studies (Garderet, Dulphy, Douay, Chalumeau, Schaeffer, Zilber, Lim, Even, Mooney, Gelin, Gluckman, Charron and Toubert, 1998, Murray et al., 2012, Raaphorst et al., 1994), nonrandom usage of TCRBV and TCRBJ genes also existed in our studies. Unlike the preferential usage of DH-proximal IGHV and DH-proximal IGHJ gene segments in the fetal B-cell repertoire(Rechavi, Lev, Lee, Simon, Yinon, Lipitz, Amariglio, Weisz, Notarangelo and Somech, 2015), no preferential usages of D-proximal or D-distal TCRBV and TCRBJ gene families were observed in the fetal TCRB repertoire in our study, indicating that their expression patterns did not correlate with their relative chromosomal position to TCRD gene segments, as was the case in adults(Lai et al., 1988).

It has been widely reported that the fetus has a shorter CDR3 length than the adult (George and Schroeder, 1992, Guo, Wang, Cao, Yang, Liu, An, Cai, Du, Wang, Qiu, Peng, Han, Ni, Tan, Jin, Yu, Wang, Wang, Wang and Ma, 2016, Rechavi and Somech, 2017, Souto-Carneiro, Sims, Girschik, Lee and Lipsky, 2005), which to some extent may limit the diversity of the CDR3 region and thus further reduce the diversity in immune cell clones. The addition and deletion of nucleotides could increase the diversity of CDR3 significantly during the formation of the junctions between gene segments, known as junctional diversity(Saada et al., 2007). Both the T-cell lineage and B-cell lineage used terminal deoxynucleotidyl transferase (TdT) to add nucleotides and probably used the same exonuclease to delete nucleotides from the coding region(Feeney, 1991). As the expression of these enzymes was developmentally regulated during ontogeny(George and Schroeder, 1992), the junctional diversity in early ontogeny was more restricted than that in adults. Consistent with previous studies (Britanova, Shugay, Merzlyak, Staroverov, Putintseva, Turchaninova, Mamedov, Pogorelyy, Bolotin, Izraelson, Davydov, Egorov, Kasatskaya, Rebrikov, Lukyanov and Chudakov, 2016, Feeney, 1991, George and Schroeder, 1992, Guo, Wang, Cao, Yang, Liu, An, Cai, Du, Wang, Qiu, Peng, Han, Ni, Tan, Jin, Yu, Wang, Wang, Wang and Ma, 2016), our results showed fewer nucleotide deletions but fewer nucleotide additions in the CDR3 of the fetus, which might lead to reduced junctional diversity and make the CDR3 sequence much closer to the germline in configuration. The dense sampling by cordocentesis across the third trimester enabled us to depict the compelling increase in CDR3 length (T1, T2 and T3), which might be in accordance with the elevated expression of related enzymes. The length of CDR3s of TCRB repertoires of cord blood in term was also longer than that in preterm blood (Le, Sper, Nielsen, Pineda, Nguyen, Lee, Boyd, MacKenzie and Sirota, 2021). For the CDR3 of the heavy chain of immunoglobulin molecule of cord blood, the length of HCDR3 in the third trimester increased with gestation weeks, and the junctional diversity could reach adult levels approximately two months after birth(Schroeder, Zhang and Philips, 2001). A similar case was also demonstrated in the intestinal TCR-delta repertoire (Holtmeier et al., 1997).

Gene usage in both BCR and TCR repertoires has been reported to be strongly influenced by genetic factors (Glanville et al., 2011, Zvyagin, Pogorelyy, Ivanova, Komech, Shugay, Bolotin, Shelenkov, Kurnosov, Staroverov, Chudakov, Lebedev and Mamedov, 2014). Twins, especially monozygotic twins who have a similar genetic identity, have been used as models to investigate the role of heredity and the environment in human immunity (Salvetti et al., 2000). Zvyagin, *et al.* found that the usage of particular TCRBV genes is strictly determined by genetic factors, while TCRBJ genes were selected randomly for recombination in monozygotic twin pairs (Zvyagin, Pogorelyy, Ivanova, Komech, Shugay, Bolotin, Shelenkov, Kurnosov, Staroverov, Chudakov, Lebedev and Mamedov, 2014). Hidetaka Tanno *et al.* further verified a more pronounced genetic influence on paired TCRαβ sequences in monozygotic twin pairs (Tanno et al., 2020). In this study, we also analysed the potential special properties of TCRB repertoires in a pedigree, including a dizygotic twin and a parental-fatal pair. We found that both the usages of TCRBV and TCRBJ genes in twins were more similar than those in other unrelated individuals, indicating that genetic factors could influence the selection of gene segments during recombination despite less common genetic identity in dizygotic twins. As the TCRBV and TCRBJ frequencies might be skewed by antigen encounter(Freeman et al., 2009), the cord blood sample used in our study could better represent the innate TCRBV and TCRBJ usage compared with adult twins. Meanwhile, no obviously closer relationship was observed in the TCRBV gene usage analysis of the maternal-fetal pair. This was in accordance with previous observations performed using mother-child pairs (Putintseva et al., 2013). Changlong Guo *et al.* also suggested that the TRB-/IGH-CDR3 repertoire of cord blood was not affected by the corresponding maternal immune status in multiple aspects(Guo, Wang, Cao, Yang, Liu, An, Cai, Du, Wang, Qiu, Peng, Han, Ni, Tan, Jin, Yu, Wang, Wang, Wang and Ma, 2016). However, there is a higher prevalence of convergent TCR-β clones between infants and mothers in preterm than that in term (Le, Sper, Nielsen, Pineda, Nguyen, Lee, Boyd, MacKenzie and Sirota, 2021). A slightly closer relevance of TCRBJ gene usage was also identified in our parental-fetal pair. Thus, more parental-fetal pair samples should be examined to confirm the results.

In summary, we comprehensively profiled the distinct characteristics of the TCRB repertoire of cord blood in late pregnancy using high-throughput TCRB enrichment sequencing. Compared to the adult control, the fetus displayed reduced repertoire diversity, a more even distribution of clones, and a shorter CDR3 length. The CDR3 length increased with fetal development. The fetus held a nonrandom usage of TCRBV and TCRBJ genes, nearly half of which were significantly different from those in adults. The differences between fetuses and adults might be attributed to the far lower exposure to antigens in utero and the ongoing maturation of the fetal immune system. These findings fill up vacancy of the features and perception about the development of fetal T cell repertoires in the third trimester of pregnancy and deepen our understanding of the health problem associated with adaptive immunity in neonates.

## Authorship

Yuan Wei, Jie Qiao and Hongmei Li developed the study conception and designs. Wei Chen and Jinmin Wang collected the study samples and the clinical information. Yan Dong carried out the experiments and analysed the data with assistance from Yuhang Cai and Xiaolei Wu. Yan Dong, Wei Chen and Jinmin Wang drafted the manuscript. Yingxin Han, Yangyu Zhao and Yuqi Wang were helpful for the discussion of this study. All the authors participated in the interpretation of the results and critically revised and approved the final manuscript.

## Acknowledgements

We thank all of the patients and volunteers who participated in this study.

## Data availability

The raw sequencing datasets in the present research are deposited in the National Genomics Data Center database and China National GeneBank DataBase; and can be obtained upon request for researchers who meet the criteria for access to confidential data.

## Supporting Information

**Figure S1.**
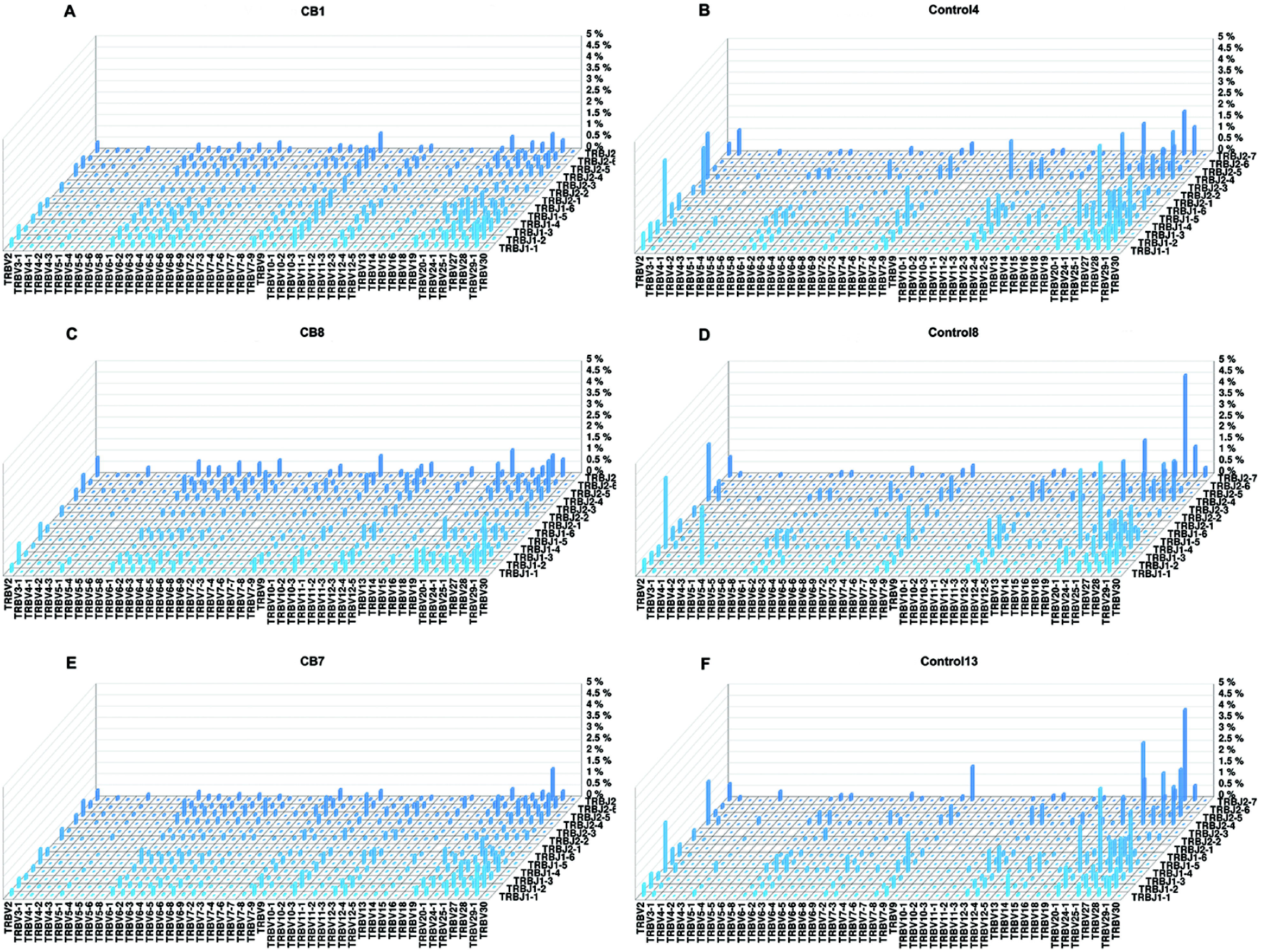
TCRB repertoire diversity in the VJ pairing pattern (3D format). (**A-F**) Visual assessment of the global TCR diversity in the VJ pairing pattern is presented. We selected one sample from each team as a representative, CB1, CB8 and CB7 (**A**, **C**, **E**), and three control samples (**B**, **D**, **F**).

**Figure S2.**
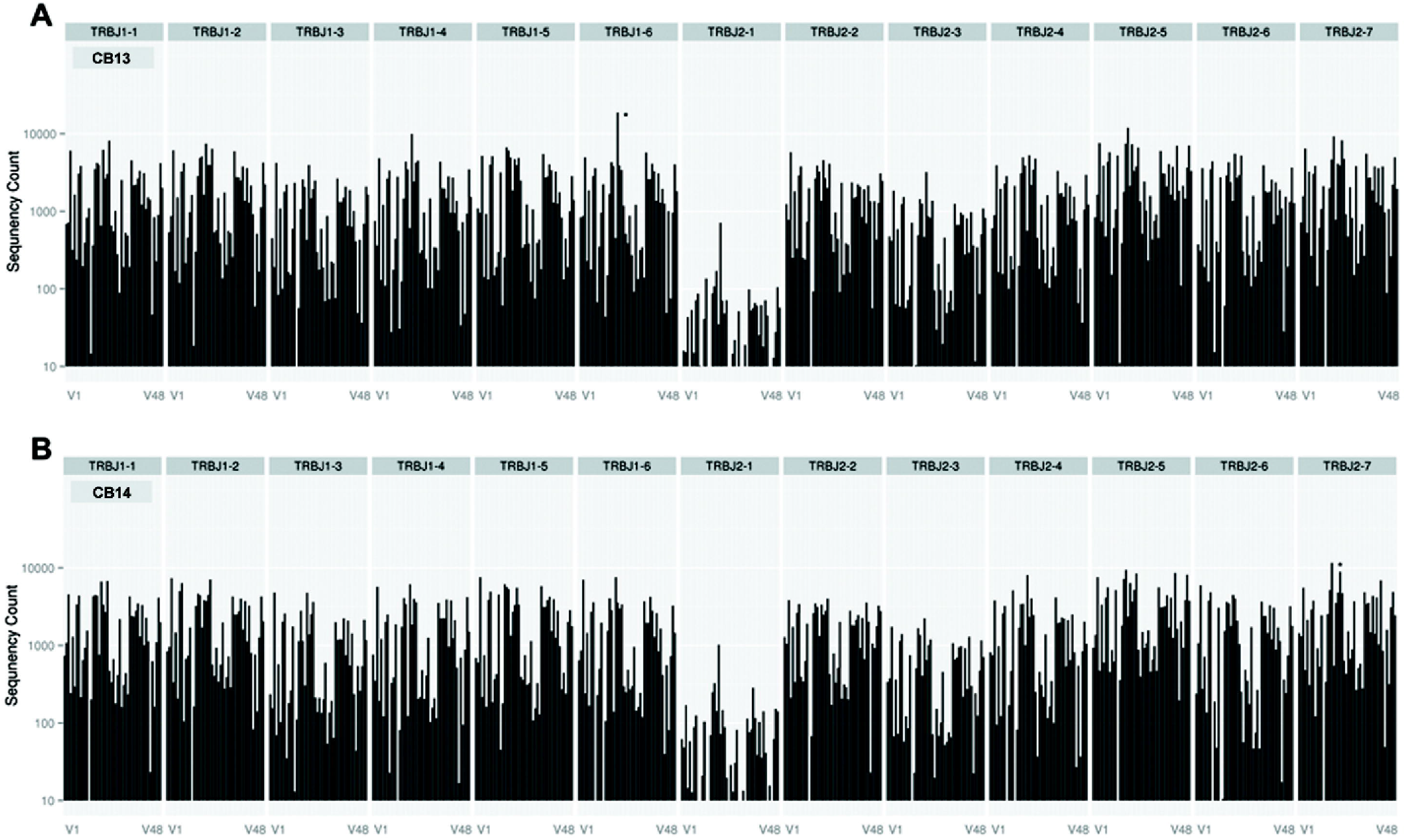
Profile of TCRBV and TCRBJ gene usage in dizygotic twin fetuses. (**A-B**) TCRBV and TCRBJ gene usage in a dizygotic twin fetus is presented. The number of CDR3 sequences belonging to a specific V gene and J gene family was calculated. (**A**) CB13; (**B**) CB14.

